# Clustering de Novo by Gene of Long Reads from Transcriptomics Data

**DOI:** 10.1101/170035

**Authors:** Camille Marchet, Lolita Lecompte, Corinne Da Silva, Corinne Cruaud, Jean-Marc Aury, Jacques Nicolas, Pierre Peterlongo

## Abstract

Long-read sequencing currently provides sequences of several thousand base pairs. This allows to obtain complete transcripts, which offers an unprecedented vision of the cellular transcriptome.

However the literature is lacking tools to cluster such data *de novo*, in particular for Oxford Nanopore Technologies reads, because of the inherent high error rate compared to short reads.

Our goal is to process reads from whole transcriptome sequencing data accurately and without a reference genome in order to reliably group reads coming from the same gene. This *de novo* approach is therefore particularly suitable for non-model species, but can also serve as a useful pre-processing step to improve read mapping. Our contribution is both to propose a new algorithm adapted to clustering of reads by gene and a practical and free access tool that permits to scale the complete processing of eukaryotic transcriptomes.

We sequenced a mouse RNA sample using the MinION device, this dataset is used to compare our solution to other algorithms used in the context of biological clustering. We demonstrate its is better-suited for transcriptomics long reads. When a reference is available thus mapping possible, we show that it stands as an alternative method that predicts complementary clusters.

## 1 INTRODUCTION

Massively parallel cDNA sequencing by Next Generation Sequencing (NGS) technologies (RNA-seq) has made it possible to take a big step forward in understanding the transcriptome of cells, by providing access to observations as diverse as the measurement of gene expression, the identification of alternative transcript isoforms, or the composition of different RNA populations [1]. The main drawback of RNA-seq is that the reads are usually shorter than a full-length RNA transcript. The growth of accession records in databases recently burst for transcripts obtained with short reads [2] but a laborious curation is needed to filter out false positive reconstructed variants that do not have enough support. Referred to as Third Generation Sequencing, long read sequencing technologies such as Pacific Biosciences [3] and Oxford Nanopore Technologies [4] have brought the opportunity to sequence full-length RNA molecules. In doing so, they relax the previous constraint of transcript reconstruction prior to study complete RNA transcripts [5]. The size of short reads constitutes indeed a major limitation in the process of whole transcript reconstitution, because they may not carry enough information to ensure the recovery of the full sequence. In addition, tools for *de novo* assembly of transcripts from short reads [5, 6] use heuristic approaches that do not guarantee the exact original transcripts to be retrieved. On the contrary long reads are prone to cover full-length cDNA or RNA molecules, thus they can inform about the comprehensive exon combinations present in a dataset. This gain in length is at the cost of a computationally challenging error rate (quite variable across protocols, up to more than 15%, although RNA reads generally show lower rates, around 9% or less [7, 8]).

In the last few years, studies have been increasing on the treatment of long read data generated via the Oxford Nanopore MinION, GridION or PromethION platforms, for transcriptome and full-length cDNA analysis [4, 9, 10, 11]. International projects have been launched and the WGS nanopore consortium (https://github.com/nanopore-wgs-consortium/NA12878/blob/master/RNA.md) has for example sequenced the complete human transcriptome using the MinION and GridION nanopores. Besides Human and microbial sequencing, this technology has also proved useful for the de novo assembly of a wide variety of species including for example nematodes [12], plants [13] or fishes [14]. It seems clear that the reduced cost of sequencing and the portable and real-time nature of the equipment will favour a wide diffusion of this technology in the laboratories compared to the PacBio technology [15] and many authors point out the world of opportunities offered by nanopores [16]. Variant catalogs and expression levels start to be extracted from these new resources [17, 18, 19, 20, 21], and isoform discovery was cited as a major application of nanopore reads by a recent review [22]. However, the vast majority of these works concern species with a reference. In this work we propose to support the *de novo* analysis of Oxford Nanopore Technologies (ONT) RNA long read sequencing data. We introduce a clustering method that works at the gene level, without the help of a reference. This enables to retrieve the transcripts expressed by a gene, grouped in a cluster. Such clustering may be the basis for a more comprehensive study that aims at describing alternative variants or gene expression patterns.

### 1.1 Problem statement

Within a long-read dataset, our goal is to identify for each expressed gene the associated subset of Third Generation Sequencing reads without mapping them on a reference. In order to group RNA transcripts from a given gene using these long and spurious reads, we propose a novel clustering approach. The application context of this paper is non-trivial and specific for at least three reasons: 1/ in eukaryotes, it is common that alternative spliced and transcriptional variants with various exon content (isoforms) occur for a given gene [23]. The issue is to automatically group alternative transcripts in a same cluster (Figure 1); 2/ long reads currently suffer from difficult indel errors at high rate [7, 8]; 3/ all genes are not expressed at the same level in the cell [24, 25, 26]. This leads to an heterogeneous abundance of reads for the different transcripts in presence. Thus clusters of different sizes including small ones are expected, which is a hurdle for most algorithms, including the prevalent methods based on community detection [27].

**Figure 1:**
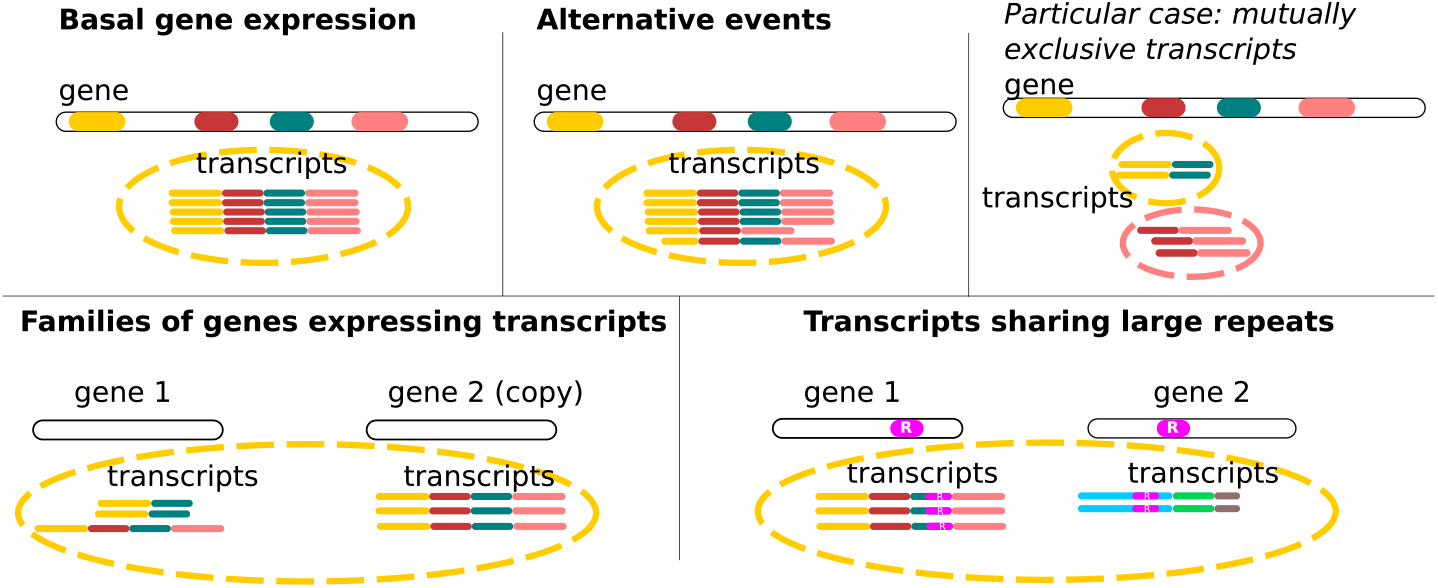
Clustering scenarii. In the case of basal gene expression and alternative events (described below), with the exception of mutually exclusive transcripts, it is expected that all transcripts of a gene will be grouped together in a single cluster. Very small exons or very long retained introns (not shown) can also be limitations according to the mapping tool strategies. In the more complex case of families of genes, two or more copies of paralogous genes can express transcripts at the same time. If these transcripts share a common exonic content and if the gene sequences have not diverged too much (to allow overlap detection), transcripts from this family of genes are clustered together, despite coming from different loci. Although this is an algorithmic limitation, it can be interesting to group these sequences together, as they likely share similar functions. A like scenario occurs for transcripts sharing genomic repeats (such as transposable elements).

Our method starts from a set of long reads and a graph of similarities between them. It performs an efficient and accurate clustering of the graph nodes to retrieve each group of gene’s expressed transcripts (detailed in section “*MATERIALS AND METHODS*”). A second contribution of our work is an implementation of the clustering algorithm in a tool dubbed CARNAC-LR (**C**lustering coefficient-based **A**cquisition of **RNA C**ommunities in **L**ong **R**eads) inserted into a pipeline (see section Results). The input of this pipeline is a whole raw reads dataset, with no prior filter or correction needed. The output is a set of clusters that groups reads by gene without the help of a reference genome or transcriptome.

### 1.2 Background

Early attempts to solve this problem can be traced back to before the age of NGS: in the NCBI Unigen database [28] Expressed Sequence Tags (ESTs) are partitioned into clusters that are very likely to represent distinct genes. In fact, clustering has been the basis for gene indexing in major gene catalogues like Unigene, HGI, STACK or the TIGR Gene Indices [29, 30]. Moreover this problem has appeared in many disciplines, taking different forms according to the application domain. Many works applied to sequence clustering put efforts in finding the most efficient way to compute similarity but remained quite basic in their clustering scheme (e.g. CD-HIT [31], SEED [32], Uclust [33], DNA-CLUST [34]). They essentially tried to avoid by these simple schemes all versus all pairwise comparison of sequences, that became a major issue with the advent of NGS and meta-transcriptomics. These approaches and the underlying similarity measures were thought for highly similar sequences, and are also popular for applications out of the scope of this paper such as clustering OTUs. In the context of proteins [35], spectral clustering has been shown to provide a meaningful clustering of families. It uses the Blast E-value as a raw distance between sequences and it integrates all of them to decide with a simple K-means clustering a global partition of protein sequences. This type of work cannot easily be extended to the comparison of reads, which are much less structured than protein sequences. To our knowledge no article has been published so far using spectral clustering on RNA reads. In the context of RNA, using Sanger reads then short reads, many approaches used simple single linkage transitive-closure algorithms (EST clustering such as [36, 37, 38]), i.e. searched for connected components in a graph of similar sequences. Single linkage clustering is often used for expression data as two similar sequences are meant to merge their clusters into a single one. A counterpart of simple search for clusters is that it can easily lead to chimeric clusters, especially because of repetitions.

Thus, more advanced clustering strategies have been developed on graphs, which use the topological properties of the graph to select relevant classes. Roughly speaking, resolution strategies can be classified into two trends according to applications and the community of affiliation: a *graph clustering strategy* based on the search for minimal cuts in these graphs and a *community finding strategy* based on the search for dense subgraphs. Our own approach aims to combine the best of both worlds. The first approach generally searches for a partition into a fixed number of clusters by deleting a minimum number of links that are supposed to be incorrect in the graph. The second approach frequently uses a modularity criterion to measure the link density and decide whether overlapping clusters exist, without *a priori* regarding the number of clusters. Given that it is difficult to decide on the right number of clusters and to form them solely on the basis of minimizing potentially erroneous links, the main findings and recent developments are based on the community finding strategy and we will focus our review on this approach. *Modularity* measures the difference between the fraction of edges within a same cluster and the fraction of edges that would be observed by chance given the degree of each node. In particular modularity-based partitioning of sequences [39] was applied for discovering protein homology [40] or repeat sequence clustering [41]. Improved state-of-the-art methods consider either overlapping communities or hierarchical communities. A well-established method for overlapping communities is the Clique Percolation Method (CPM) [42]. CPM came with applications such as identification of protein families [43, 44].

Finally recent works [45] rely on Louvain algorithm [46] that is also based on modularity and is looking for a hierarchy of clusters, by practicing a multi-level optimization that merges the clusters initially reduced to one element as long as the modularity increases. This algorithm is fast because it uses a greedy strategy and is quite popular for extracting communities from large networks. However, like the other algorithms based on modularity, it suffers from two drawbacks: it has difficulty dealing with small clusters and is unstable in that, depending on the order of application of merges, it can produce very different solutions that are difficult to compare [47].

Clustering problems applied to the specificity of long reads start to emerge. Such needs were already of concern in the past long read literature [19, 48] and are even more acute when a mapping strategy cannot be taken into consideration. We place ourselves in the particular framework of *de novo* identification. While several works based on long read mapping on a reference were proposed, methodological contributions that would enable to make the most of this promising data remain rare in particular for non model species. To our knowledge, two contributions [49, 48] propose respectively to *de novo* detect alternative variants and to cluster and detect isoforms in long reads transcriptome datasets. However these tools highly rely on the property of high accuracy proposed by Pacific Biosciences Consensus Circular Sequence (CCS) long reads, thus do not apply to ONT reads. The method we propose is much more robust to noise.

## 2 MATERIALS AND METHODS

### 2.1 Input similarity graph

We define a similarity graph as an undirected graph in which nodes are reads and there is an edge between two nodes if the computed similarity between these nodes exceeds a fixed threshold. In such a graph, reads from a same gene are expected to be connected with one another because they are likely to share exons. In the ideal case, all reads from a gene are connected with one another. It is therefore a clique. However, the spurious nature of data imposes the use of heuristics to detect read overlaps.

This, in addition to the presence of genomic repeats leads to the expectation of a graph with both missing edges (connection missed during the search of overlapping reads) and spurious edges (wrong connections between unrelated reads), which motivates the development of tailored clustering methods.

### 2.2 Clustering long reads

#### 2.2.1 Clustering issue and sketch of the algorithm

##### Problem formalization

In the following, we describe the clustering algorithm that is the main contribution of this paper. Our method makes no assumption on the number of expressed genes (i.e. clusters/communities), nor on the size distribution of such communities, and it needs no input parameters. As we want to realize a partition of the graph, there are no intersecting communities (no read belongs to several gene families) and every node belongs to a community (each read is assigned to a gene). As mentioned previously, the expected subgraph signature of a gene in the graph of reads is a community, that is, a cluster of similar reads. Clustering seeks to maximize intra-cluster similarity and minimize inter-cluster similarity. To measure the density of a connected component, we use the clustering coefficient (*ClCo*) [50] rather than modularity. Indeed, we assume that a gene should be represented by a complete subgraph (clique) in a perfect similarity graph. The value of *ClCo* measures the concentration of triangles in a given subgraph (see section “Selection of community founding node”), and this coefficient is more directly connected to the notion of clique than modularity. The paragraph “*Results on theoretical instances*” in the Results section clearly demonstrates the advantage of *ClCo* on this aspect over modularity.

Although we have designed a parameter-free method, its foundation is a problem depending on two parameters, the number *k* of clusters and the cutoff *θ* on the *ClCo* value. Specifically, the original problem is formalized as follows:

###### Definition 1

*A community is a connected component in the graph of similarity having a clustering coefficient above a fixed cutoff θ. An optimal clustering in k communities is a minimal k-cut, that is, a partition of the graph nodes in k subsets, that minimizes the total number of edges between two different subsets (the cut-set)*.

We assume that the overlap detection procedure (section “*First step: computing similarity between long reads*”) has good specificity (it does not produce a lot of false positives). This can be ensured by carefully tuning the parameters of this procedure. The logic behind the search for a minimum cut in the graph is that most of the edges of the initial graph should therefore be kept during clustering. This problem is known to be NP-hard for *k* ≥ 3 [51]. Another source of complexity is that we don’t know in advance the number of communities, so we have to guess the value of *k*. One should thus compute the *k*-cut for each possible value between 1 and the maximum, which is the number of reads. Solving this problem is not feasible for the large number of reads that have to be managed. We are thus looking for an approximation of the solution by using an efficient heuristic approach exploring a restricted space of values for *k*. Finally, the second parameter, the cutoff *θ*, is not known either. The algorithm has thus to loop over all possible values, that is, all *ClCo* values for a given connected component. In practice it is sufficient to sample a restricted space of possible *k* values.

##### Algorithm overview

Shortly, our community detection algorithm is composed of two main steps. The first one looks for an upper bound of the number of clusters *k*. To this aim, we relax the condition of disjoint communities and look initially for star subgraphs (a read connected to all reads similar to it) having a clustering coefficient above a certain cutoff. This corresponds to detecting well-connected reads, called seed reads, using *ClCo* and node degrees (detailed in section “*Selection of community founding nodes*”). They form the basis of communities with their neighborhood.

The main challenge is then to refine the boundaries of each community (section “*Refinement of community boundaries*”) in order to fulfill the partition condition. During this process, the value of *k* is progressively refined by possibly merging clusters whose combination produces a better community (greater *ClCo* value). The other possibility of refinement is to assign nodes to a community and remove them from another. If *x* edges between a node and its previous community are removed, the cut size of the partition is increased by *x*. This core algorithm is run for different cutoff values to obtain different partitions that we compare. We keep the partition that is associated to the minimal cut (i.e. number of edges removed when computing this partition). The pseudocode of the implemented algorithm is given in the “Supplementary material”. In the following, the different steps of the implementation are detailed.

#### 2.2.2 Generation of partitions

In order to generate and compare different partitions for the graph, we define cutoffs that rule the generation and refinement of communities. The cutoffs can be seen as the level of connectivity at which a community can be generated ((a,b) steps and (c) merge step in Figure 2). In the basic algorithm, for each connected component, all *C_l_C_o_* are computed in the first place, and partitions are built for each non-zero *ClCo* value as a cutoff. In the end, only one partition is retained, associated to the minimal cut (step (d) in Figure 2). However we have reduced the number of possible cutoff values for the sake of scalability (see section Implementation choices). In the following, each step is described for a given cutoff value.

**Figure 2:**
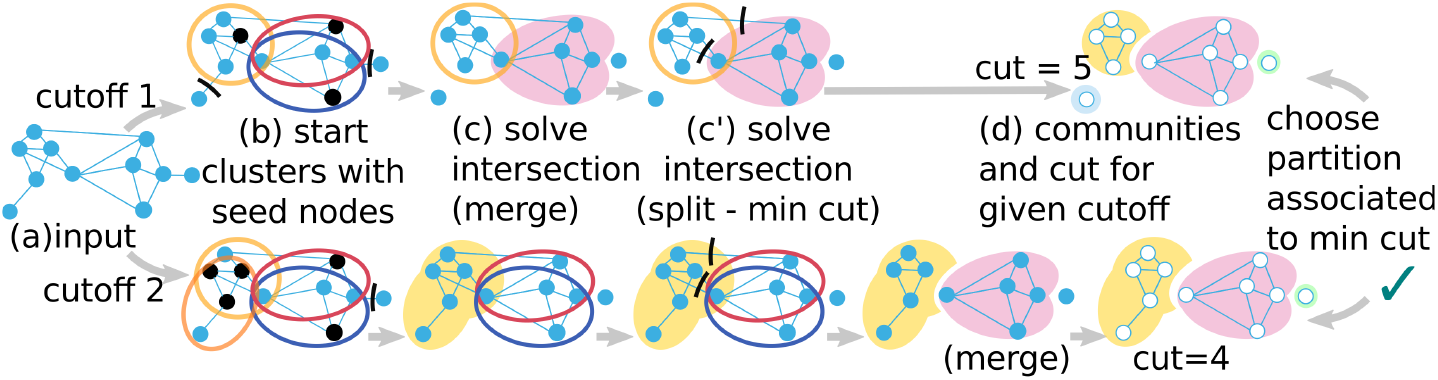
Summary of the algorithm. (a) All *ClCo* and degrees are computed. Each *ClCo* value is a cutoff. For a given cutoff, (b) different cutoffs yield different seed nodes (black stroke) that initiate clusters with their neighborhood (section “*Selection of community founding nodes*”). (c, c’) Boundaries of each cluster are then refined. Intersection between clusters are solved either by (c) merging them or by (c’) splitting (section “*Refinement of community boundaries*”). (d) The communities at different cutoffs evolve in different partitions. In the end we keep only the best partition according to our criterion, i.e. minimizing the cut.

#### 2.2.3 Selection of community founding nodes

Let 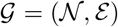 be an undirected graph of reads. Let *n*_*i*_ be a node (read) from 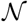 and 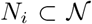 its direct neighborhood. Let *deg*(*n*_*i*_) be the number of edges connecting *n*_*i*_ to its direct neighbors (similar reads), i.e. *deg*(*n*_*i*_) = |*N*_*i*_|. For each node *n_i_ ∈ N* with degree *deg*(*n*_*i*_) *>* 1, we first compute the *local clustering coefficient*:

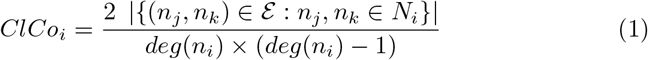

Nodes of degree 0 and 1 have a *ClCo* of 1. This local coefficient represents the *cliqueness* of the *N*_*i*_ ∪ *n*_*i*_ set of nodes. The closer to 1, the more the set of nodes is inter-connected, which witnesses a group a reads that potentially come from the same gene. By contrast, the subgraph induced by a node with a *ClCo* of 0 and its neighbours is a star (i.e. a tree whose leaves are all the neighbours). If the coefficient is close to 0, the nodes are weakly connected and are unlikely to come from the same gene. In order to prevent unwanted star patterns we add an auxiliary condition for nodes to be eligible seeds, described in “Supplementary Material”. At this point it is possible that two or more communities intersect.

#### 2.2.4 Refinement of community boundaries

Community refinement aims at solving the conflicts of intersecting communities. Communities intersection happen because of spurious connections in the graph due to the creation of edges between unrelated reads in the first step.

The intersecting communities are looked up pairwise in order to assign nodes of the intersection to only one community. In fact two cases have to be distinguished. Either the edges between two communities are estimated spurious and these communities must be seen separated (*split*, (c’) step in Figure 2 (the pseudocode for the *split* procedure is also given in the “Supplementary material”), or the edges have sufficient support and the two communities have to be merged to obtain the full gene expression (*merge*, (c) step in Figure 2). In order to decide between the two, we use again the *cliqueness* notion. This time we introduce an *aggregated clustering coefficient* of the union of two nodes *n*_*i*_ and *n*_*j*_:

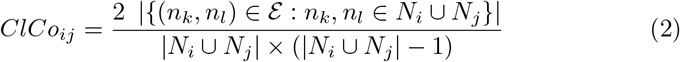

If the value of *ClCo*_*ij*_ is greater than or equal to the current cutoff, we consider that there is a gain in connectivity when looking at the union of the two communities and they are merged. In the other case, the nodes of the intersection are reported to only one of the two communities. We remove the edges connecting these nodes to one or the other cluster according to which realizes the minimal cut. In case of ties for the cut, the algorithm uses a second criterion described in the “Supplementary material”. The global result depends on the order in which pairs of clusters are compared. This order is carefully designed. First the communities associated to the two nodes of greatest degree (and secondly maximal *ClCo*) are chosen, the intersection is resolved and the first community is updated. Then, it is compared to the third best community that intersected it if it exists, and so on until all intersections are solved. This way, we start the comparison by the most promising communities that combine reliability (they are well-connected subgraphs) with a high potential of resolution (they likely are the biggest communities, thus solve intersections for many nodes). On the contrary, communities associated to small subgraphs and relatively low *ClCo* are only resolved afterwards.

#### 2.2.5 Complexity and Implementation choices

Our algorithm has a quadratic component to compare sets to get clusters. In addition, it explores the whole space of clustering coefficients to fix cutoffs. It results in a time complexity that could theoretically be cubic in the number of reads at worst, which is incompatible with processing large datasets.

In order to cope with noise in the input graph, we introduce features to simplify the graph (disconnect loosely connected nodes) and to control the space of research of the possible partitions. In practice these features are also key to reduce the complexity of our approach. Our experiments showed that the running time is reasonable, clustering millions of reads in a few hours. Two key ideas to obtain this result have been to reduce the number of cutoffs and to disconnect the articulation points [52] to reduce the size of connected components in the graph. Details are given in the “Supplementary material”.

Indeed, the most costly phase relies on the treatment of the largest connected components. In these components, many clustering coefficients values are very close and their variation is mainly an effect of noise. Introducing a rounding factor when computing the *ClCo* results in a neat decrease in the number of different values observed, which drastically limits the number of iterations required for the main loop, while providing a very good approximation of the minimal cut. In addition, an upper bound is set on the number of sampled values (100 by default).

We also chose to disconnect the *articulation points* of the graph to remove nodes that can be targeted as potential bridges between two correct clusters. These are nodes whose removal increases the number of connected components in the graph. Such nodes can be spotted as problematic as we do not expect a single read to be the only link between many others. Their detection can be done with a DFS in linear time for the whole graph.

Our algorithm has been also carefully designed with respect to the features of long read clustering. To obtain a *O*(*n.log*(*n*)) complexity with respect to the number *n* of reads, we have made the following assumption: The degree of each node is bounded by a constant, i.e. there is a limited number of transcripts that share similar exons. This ensures that the clustering coefficient of all nodes is calculated in linear time. The most complex operation is the initial sorting of nodes, first by decreasing degree value, then by decreasing *ClCo* value, which can be achieved in *O*(*n.log*(*n*)). Moreover, since each cluster is initially built on a seed read (see paragraph Selection of community founding nodes), it intersects with a bounded number of clusters. Since the loop for making a partition from overlapping clusters is based on a scan of intersections, it is achieved in linear time wrt the number of reads.

### 2.3 Validation procedure

#### 2.3.1 Production of validation material

##### RNA MinION sequencing

cDNA were prepared from 4 aliquots (250ng each) of mouse commercial total RNA (brain, Clontech, Cat# 636601 and 636603), according to the Oxford Nanopore Technologies (UK) protocol “1D cDNA by ligation (SQK-LSK108)”. The data generated by MinION software (MinKNOWN, Metrichor) were stored and organized using a Hierarchical Data Format. FASTA reads were extracted from MinION HDF files using poretools [53]. We obtained 1,256,967 nanopore 1D reads representing around 2Gb of data with an average size of 1650bp and a N50 of 1885bp.

##### Mapping to obtain reference clusters for validation

We compute “ground truth” clusters for validation purpose, using a sensitive third-party protocol based on mapping on a reference. Nanopore reads from the mouse brain transcriptome were aligned to the masked mouse genome assembly (version GRCm38) using BLAT [54]. For each read, the best matches based on BLAT score (with an identity percent greater than 90%) were selected. Then, those matches were realigned on the unmasked version of the genome using Est2genome [55] that is dedicated on precise spliced-mapping on reference genome. Reads that corresponded to mitochondrial and ribosomal sequences were discarded. Next, Nanopore reads were clustered according to their genomic positions: two reads were added in a given cluster if sharing at least 10nt in their exonic regions. For the whole data experiment, all reads that could be mapped on the reference were taken into account (501,787). Due to repeats (paralogy, transposable elements…), some reads were mapped at multiple loci on the reference. When a given read maps on several loci, such loci are gathered in a single expected cluster (12,596 expected clusters). This means that for instance reads from copies of paralog genes that have not diverged to much or reads containing a copy of a transposable elements are expected to be in the same cluster.

#### 2.3.2 Clusters’ goodness assessment metrics

To assess the results, we used recall and precision measures, which are standard measures to assess the relevance of biological sequence clustering [56]. The recall and precision measures are based on reference clusters obtained by mapping for this validation. They are computed based on representative graphs [57]. These measures were already used to assess the relevance of biological sequence clustering [56]. Let 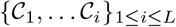 be the set of clusters found by the clustering method tested, where *L* is the number of predicted clusters. Let 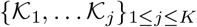 be the set of “ground truth” clusters, where *K* the number of expected clusters. Let *R*_*ij*_ be the number of nodes from 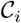 that are in “ground truth” cluster 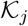. We compute the recall and the precision such as:

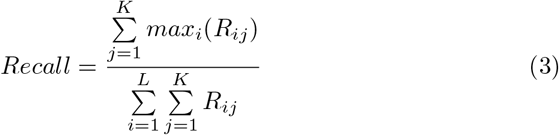

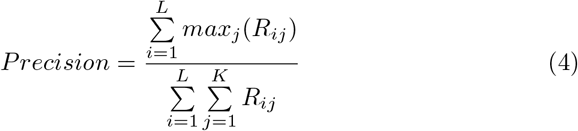

Note that the “ground truth” is not really available and that it is estimated from mapping results on the reference genome. The recall expresses the mean over all clusters of the fraction of relevant reads in a result cluster out of the expected read population of this cluster. It presents to which extent cluster are complete. The precision expresses the mean over all clusters of the fraction of relevant reads among the population of a result cluster. It shows the clusters’ purity. The F-measure is a summary measure computed as the weighted harmonic mean between precision and recall. Recall and precision are tailored to express the confidence that can be placed in the method, according to its ability to retrieve information and to be precise. We complementary assess the closeness of the computed clusters as compared to mapping approaches. Let *χ*_0_ be the reference partition (set of clusters obtained by mapping), and *χ* the partition obtained using a given clustering method. Then *a*_11_ is the number of pairs of nodes that are placed in a same cluster in *χ*_0_ and *χ*_1_. *a*_00_ indicates the number of pairs for which nodes are placed in different clusters both in *χ*_0_ and *χ*_1_. *a*_10_ (resp. *a*_01_) is the number of pairs of nodes placed in the same cluster in the reference *χ*_0_ (resp. *χ*) but in different clusters in (resp. _0_). Based on those, a metric such as the Jaccard index shows the match between the reference and computed partitions:

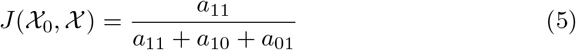

The Jaccard index is between 0 and 1. The closer to 1, the more the set of clusters computed by a method is close to the “ground truth” set of clusters predicted.

## 3 Results

All experiments were run on Linux distribution with 24 Intel Xeon 2.5 GHz processors, 40 threads and 200 GB RAM available. First we present the tool we have developed and made available for large scale long-reads clustering. We demonstrate it performs well on a canonical example on which other clustering approaches were assessed (section “*Results on theoretical instances*)”. We compare our approach to well established community detection methods and demonstrate its relevance on long read application. Then we validate our method’s results by comparing them with independent clusters obtained by mapping on a real size dataset. In these two parts (sections “*Comparison to state of the art clustering algorithms*” and “*Biological relevance*”), reads from the brain mouse transcriptome were used in order to access a “ground truth” via a reference. Then we show that our approach can offer an alternative to the classical mapping approach even when a reference is available.

### 3.1 CARNAC-LR, a software for long read clustering

#### 3.1.1 Input/Output

We implemented our novel algorithm presented in section “*MATERIALS AND METHODS*” in a tool called CARNAC-LR. CARNAC-LR comes with a pipeline. It starts with the computation of long read similarities by a program called Minimap [58] and then produces the clusters using CARNAC-LR. The pipeline’s input is a FASTA file of reads. The output is a text file with one line per cluster, each cluster containing the read indexes. Each read is represented by its index in the original FASTA file during CARNAC-LR computation. Then each cluster can easily be converted to a FASTA file, where using indexes, each read’s sequence is retrieved from the original file (a script is proposed for doing this task).

#### 3.1.2 First step: computing similarity between long reads

We chose the tool Minimap for its efficiency and its very high level of precision on ONT and PB [59], among other recent methods that can compute similarity or overlaps between long reads despite their error rate [60, 61, 62, 63]. To generate the similarity graph for CARNAC-LR, Minimap version 0.2 was launched with parameters tuned to improve recall (`-Sw2 -L100 -t10`). It produces a file of read overlaps in .paf format.

#### 3.1.3 Second step: clustering

Minimap’s output is converted into a graph of similarity file, where a node represents a read and an edge a sequence similarity between two reads above a certain threshold (see [58]). Such graph is then passed to CARNAC-LR that retrieves and outputs the gene clusters. CARNAC-LR benefits from parallelization. A thread can be assigned to the treatment of a single connected component, thus many connected component can be computed in parallel. Further results on scalability are provided in the “Supplementary material”.

### 3.2 Method validation

#### 3.2.1 Results on theoretical instances

Fortunato et al. [64] proposed to test the resolution limit of community detection on a ring of 30 cliques of 5 nodes interconnected through single links. The Louvain algorithm finds the partition in cliques at the first level of the hierarchy and build groups of 2 cliques at the second and last level. CARNAC-LR finds the correct partition in cliques without its articulation node filter. As this instance was easy to retrieve, we slightly complicated the initial example: we made the cliques bigger (size 7) and cliques are interconnected through two links on different nodes (see Figure 3, left). We provide an example of the resolution achieved by Louvain’s algorithm on this new problem to illustrate its difficulty (see Figure 3, right). It cannot find the partition in cliques and moreover, the cliques are not always split at the same place. Contrary to modularity-based approaches [64, 46] CARNAC-LR successfully reported the 30 expected clusters of cliques (Figure 4).

**Figure 3:**
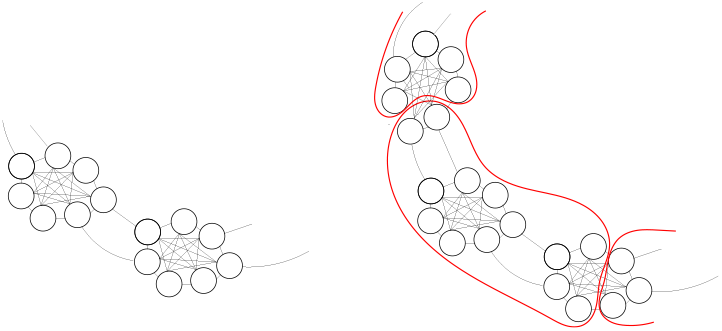
On the left we provide an example to show how 7-cliques are connected in our example design. In total, a ring of 30 7-cliques is used. On the right we illustrate Louvain’s result on this instance. The clusters formed by Louvain are in red, spanning several cliques.

**Figure 4:**
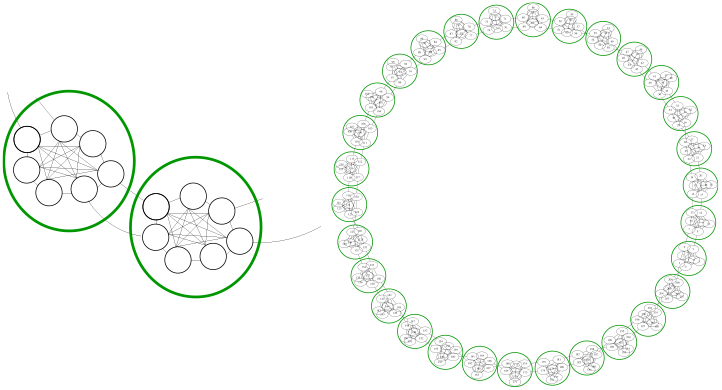
CARNAC-LR result clusters on the ring of 7-cliques. A close-up on two cliques is shown on the left. Each cluster output by CARNAC-LR is represented by a green circle. It can be seen that cliques are separated from each other as our method identifies them as independent clusters.

#### 3.2.2 Comparison to state of the art clustering algorithms

We show results of state of the art algorithms and compare them to our tool’s results. For scaling purpose, we chose to perform the benchmark on a subset of 10K reads (10,183 mouse reads within 207 reference clusters determined by mapping, section “*Production of validation material*”). Such sampling can accentuate the low expression effect in the subset. We have thus checked on a second 10K sample from chromosome 1 only to also account for highly expressed genes that results have the same trend than those presented (shown in “Supplementary material”). We compared CARNAC-LR results to algorithms we identified as close to the solution we propose. We evaluated four state-of-the-art methods that were previously applied to similar problems of biological clustering: single linkage transitive-closure [36, 37, 38], *modularity* [40, 65, 66], Clique Percolation Method [43, 44] and Louvain [45, 67]. Results are presented in Table 3.2.2. Our method has the best precision and the best overall trade-off between precision and recall as shown by the F-measure. It also has the highest Jaccard index among all tested approaches. The transitive closure approach suffers from low precision. The modularity-based approach fails to find good clusters for this graph, with both low recall and precision. CPM was tested with values for input parameter *k* ranging from 3 to 50 (no community found for greater values). Results are presented for *k*=5 and *k*=50 and summarize the behavior of this approach on our input graph. For low values of *k*, CPM outputs more clusters and has better recall than for high values. However its precision is globally low. For higher values of *k*, the results are strongly enhanced but represent only a small fraction of the input. Louvain’s results are presented for the last iteration of their algorithm. We also tested results after the first iteration, with similar trends. Despite showing the best recall, Louvain’s precision is too low to reach a high F-measure or Jaccard index. As CARNAC-LR is conceived for general pipelines making the complete analysis of gene variants, it is important that is does not mix two unrelated genes in a same cluster. Thus our approach is more conservative than CPM, and it shows comparatively good results in any case, and furthermore needs no input parameter.

**Table 1:** Comparison with state of the art methods. The benchmark was realized on a 10K reads dataset from the mouse brain transcriptome. Recall precision and Jaccard Index are presented (see Equation 5, 3 and 4) to assess for the goodness of communities detection. CPM5 (resp. CPM50) designates the CPM algorithm using *k* = 5 (resp. *k* = 50).

|  | Recall (%) | Precision (%) | F-measure (%) | Jaccard index |
| --- | --- | --- | --- | --- |
| Transitive closure | 75.74 | 5.614 | 13.62 | $7.3E^{-4}$ |
| Modularity | 60.70 | 71.16 | 65.51 | $9.7E^{-2}$ |
| CPM5 | 79.00 | 69.35 | 73.86 | $3.5E^{-1}$ |
| CPM50 | 49.21 | 89.92 | 63.60 | $7.6E^{-2}$ |
| Louvain | **88.58** | 14.91 | 25.53 | $1.1E^{-3}$ |
| CARNAC-LR | 65.0 | **98.41** | **86.62** | **$7.9 E^{-1}$** |

#### 3.2.3 Comparison to other nucleic sequence clustering tools

We have just situated the CARNAC-LR algorithm in relation to existing general cluster detection methods, but we still have to compare our pipeline to other tools dedicated to the comparison of nucleic sequences that have been developed for the same clustering task. We started from one of the most powerful tools currently available, Starcode [68], which was designed for reads correction and offers a benchmark that we have adopted of the most widely used clustering tools. It includes CD-HIT [31], SEED [32] and Rainbow [69]. We emphasize that none of these tools have been designed to work with ONT reads. Created before the full development of long reads technology, they have not surprisingly proved completely ill-suited to clustering these long reads. For this test, we used the same mouse dataset as in the previous section. The methods stumble on two data features, the error rate and the length of the sequences. SEED for instance is designed to create clusters with sequences that differ from at most 3 mismatches, thus finds no clusters. Starcode is not adapted to the size of ONT sequences and terminates with an error message. We have tried to increase the maximum size allowed for sequences (initially set at 1024) but the memory consumed is growing rapidly and reasonable capacities (200GB) are quickly exceeded. We then tried to perform the calculation by rejecting the longest reads but as well as SEEDS, Starcode authorizes a limited distance between pairs of sequences (a maximum Levenshtein distance of 8) which is far too small for ONT reads, resulting into singleton clusters. Rainbow only accepts paired reads such as those sequenced in RAD-seq short reads experiments and cannot be adapted to our problem. Finally the most flexible tool, CD-HIT, was the sole to give results. It has been used in its “EST” version. We tested different values for sequence identity threshold (`-c`), that can be decreased down to 0.8. We report only the best result, reached for `-c` 0.8. It is far below the result obtained by CARNAC-LR (F-measure up to 41.96% due to low recall, against 86.62% for CARNAC-LR). In addition, our pipeline is substantially faster with memory consumption in the same range (within a factor of 2). In view of these results, we added to the benchmark the only other *de novo* clustering tool that, to our knowledge, is designed to work with long reads, Tofu [49]. Unfortunately, Tofu highly relies on the specificity of Pacific Bioscience RNA protocol (Isoseq) sequences, and cannot be run with ONT reads. Incidentally, the aim of Tofu differs from CARNAC-LR as it is expected to retrieve one cluster per isoform rather than one cluster per expressed gene. A detailed summary of this benchmark result is presented in “Supplementary materials”. Again, another sampling on mouse chromosome 1 was used to perform a second benchmark that presents same conclusions, as also shown in “Supplementary material”.

### 3.3 Biological relevance

#### 3.3.1 Validation on a real size dataset

##### Clusters goodness

In this experiment we demonstrate the quality of *de novo* clusters obtained by CARNAC-LR. We used the subset of reads that could be mapped to the mouse genome reference (501,787 reads) as a way of comparison to assess the biological relevance of our clusters. CARNAC-LR’s results were computed using 43GB RAM and 18 minutes.

The mean recall for CARNAC-LR was of 75.38% and the mean precision was 79.62%. In other words, retrieved clusters are 75.38% complete on average, and an average 79.62% portion of the clusters is composed of unmixed reads from the same gene. In order to evaluate if our method’s recall and precision is consistent independently of the genes’ expression levels, we computed expression bins. For a given gene, we use the number of reads of the “ground truth” cluster to approximate an expression. Any “ground truth” cluster with 5 or less reads is placed in the first bin, and so on for 5-10, 10-50 and ≥ 50 reads categories. Each of the four bin represents quartiles of expression, which means there is an equal number of clusters in each bin. Figure 5 presents the recalls obtained for binned expression levels and shows our approach’s recall and precision remain consistent despite the heterogeneous coverage in reads.

**Figure 5:**
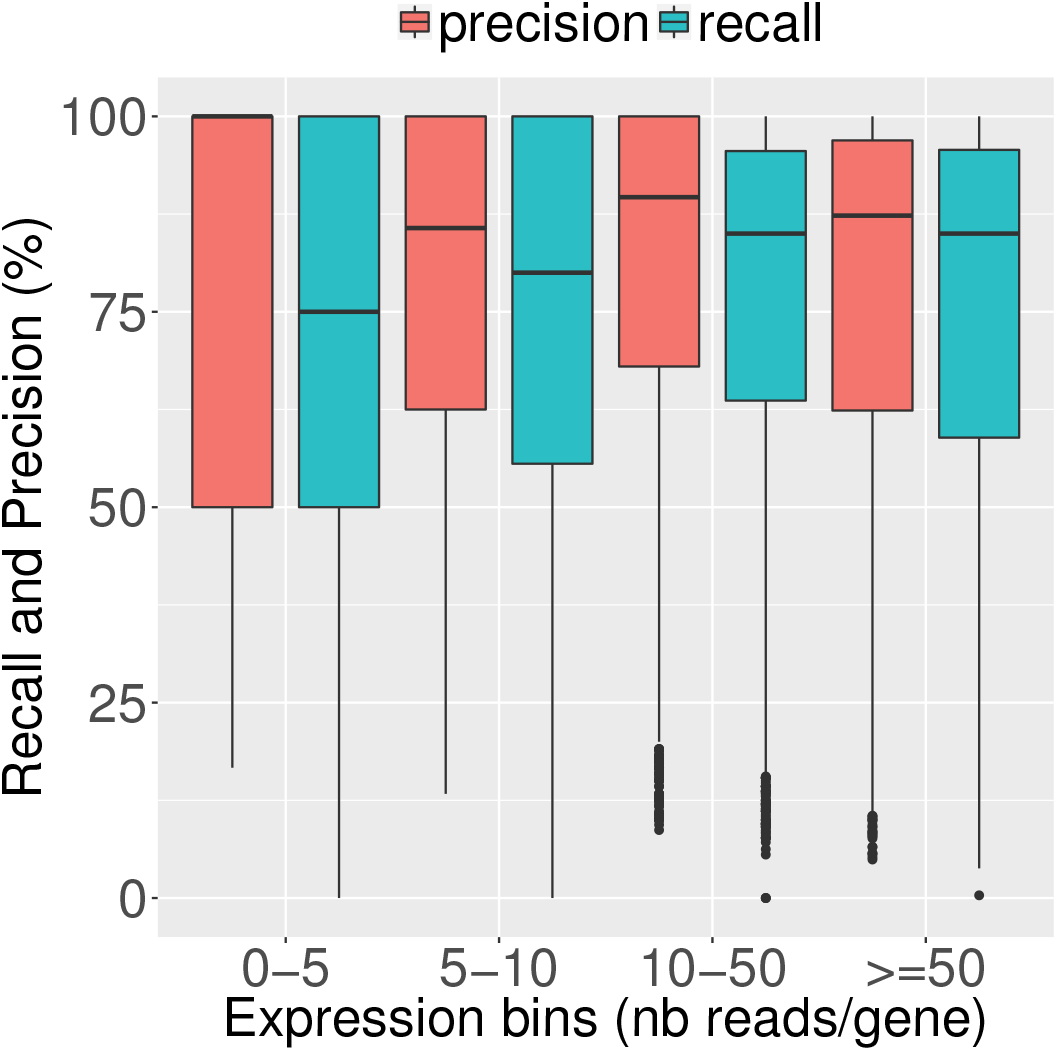
Assessed mean recall and precision of CARNAC-LR+Minimap. They were computed on mouse reads using clusters found by mapping on a reference as a “ground truth” (see Equations 3 and 4). Expression bins are computed from quartiles of expression predicted by mapping and represent the number of mapped reads by gene. Mean precision and recall over all clusters falling in theses bins were then calculated.

Furthermore, we can deduce from this plot that the numerous rather small clusters do not bias the presented mean recall and precision as even for big clusters that are more prone to loose a few reads, (i.e. ≥ 50 expression bin) these metrics remain high.

##### Output excerpt

Once CARNAC-LR is run, one can extract FASTA files for each cluster. We selected the sequences contained in a cluster after CARNAC-LR’s pass on the mouse transcriptome. In order to present a visual example of the output, we used a genome browser to display reads grouped by our approach (Figure 6). We have selected a cluster of sufficient size to be able to present a variety of isoforms. It corresponds to a gene for which mapping retrieved 120 reads. In this example, our approach retrieved 93% of the predicted gene’s reads in while including no unrelated read in the cluster. Two types of missed reads can be distinguished: 1) A group of black reads that share no genomic sequence with the majority of the gene’s transcript, because they come from an intronic region. These reads could not be linked to the others by Minimap, thus are never clustered with them, as shown in the particular case described in Figure 1. 2) Two other reads for which local connectivity was not detected by Minimap were discarded from the cluster. The plot shows different exon usage in transcripts, which reveals alternative splicing in this cluster. Thus different alternative isoforms were gathered in a single cluster as expected (see Figure 1).

**Figure 6:**
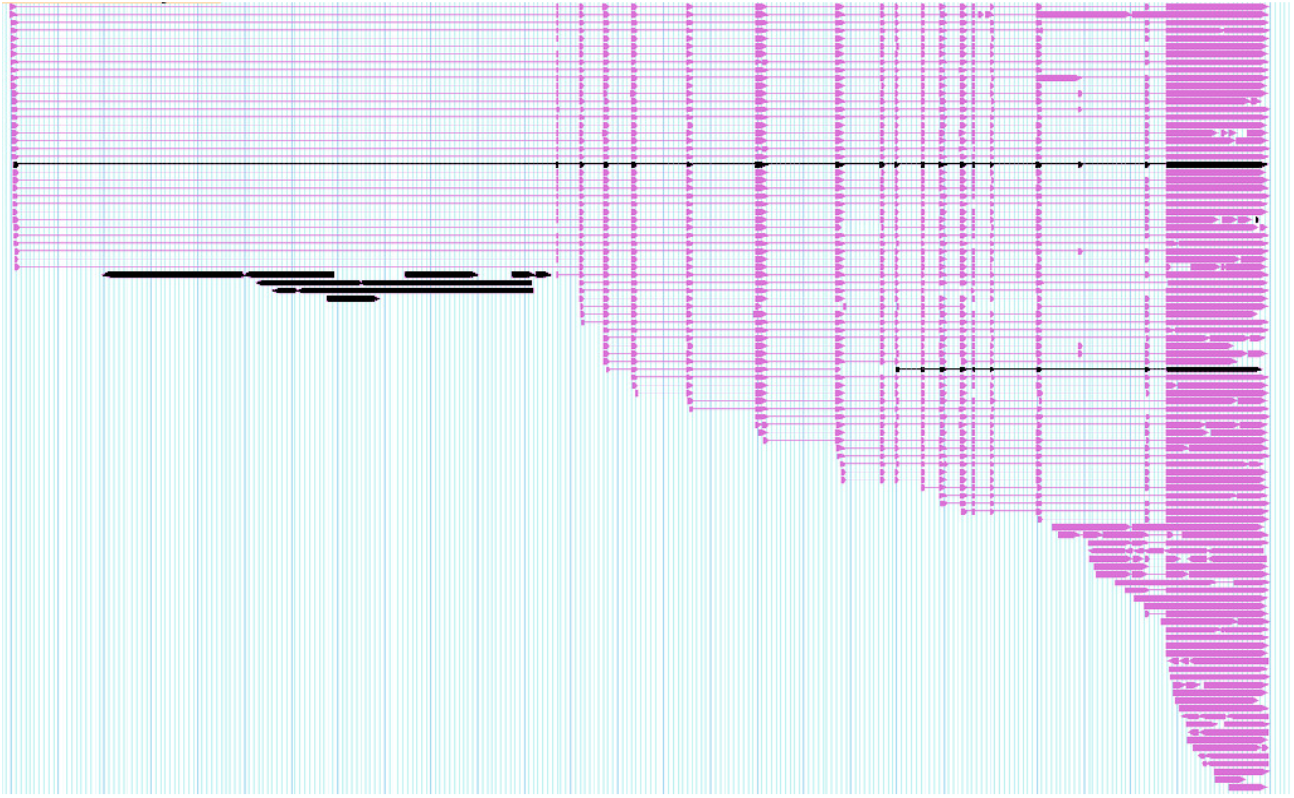
Example of CARNAC-LR’s output cluster in mouse. The output of CARNAC-LR is a text file with one line per cluster, each cluster containing the read indexes. We selected an example of 112 reads (purple) from a cluster output by CARNAC-LR. For validation purpose these reads were mapped with BLAST on an in-house igv [70] version for mouse genome. Reads are spliced-mapped, bold parts are the mapped sequences from the reads and thin parts represents the gaps between the different mapped parts of the reads. Despite the staircase effect observed in the data, this allows to notice that several types of variants were gathered. They could all be assigned to gene Pip5k1c (chr 10:81,293,181-81,319,812), which shows no false positive was present in this cluster. 8 reads (black) present in the data are missed in this cluster. The group of 6 black reads on the left represent intronic sequences and share no sequence similarity with the others and thus could appear in the same cluster.

#### 3.3.2 Complementary of *de novo* and reference-based approaches

##### Intersection and difference with the set of mapping clusters

As it does not rely on any reference information, our approach putatively yields different results than classical mapping approaches. In this section, to demonstrate the interest of CARNAC-LR even if a reference is available, we investigate the differences between the two approaches. We ran it on the full mouse brain transcriptome dataset (1,256,967 reads). We compared the intersection and difference of results of our approach and mapping. CARNAC-LR+Minimap pipeline took less than three hours (using 40 threads). In comparison, the “ground truth” clusters took 15 days to be computed (using up to 40 threads). Our approach was able to place 67,422 additional reads that were absent in the mapping procedure. It resulted into 39,662 clusters. These clusters fall in two categories (i) *common clusters* with a mix of reads treated by our approach and/or processed by mapping, or (ii) *novel clusters* that contain reads treated by our approach or mapping exclusively. Each approach performed differently on these categories.

##### Common clusters

For category (i), mapping complemented many common clusters with small amounts of reads left aside by our approach. As some reads are processed by mapping, a recall and precision can still be calculated using mapping as ground truth. We computed recall and precision based on the read fraction of clusters that could be compared with mapping. They are quite similar compared to the values obtained in the previous section (75.26% and 79.30%). This demonstrates that CARNAC-LR efficiently used the supplementary connectivity information despite the addition of potentially noisy reads.

##### Novel clusters

Conversely CARNAC-LR shows a better ability to group reads unprocessed by mapping into novel clusters (Figure 7). CARNAC-LR output 824 novel clusters (17,189 reads) of category (ii) containing at least 5 reads. In order to evaluate the relevance of these novel clusters, we remapped reads *a posteriori*, separately for each cluster, on the reference genome using a sensible approach (GMAP [71] version 2015-09-29). This operation took approximately 10 hours (using 4 threads). 19.68% of mapped reads were assigned to the MT chromosome, then chromosome 11 represented 10.85% of the reads, and other chromosomes each less than 10% of mapped reads. A third of the reads were multi-mapped (36.7%). However, on average, for each cluster 98.89% of the reads shared a common genomic locus. This is consistent with the expected results of the clustering for reads containing repeats or paralog regions (Figure 1). Finally, 5.7% of the clusters contained exclusively reads mapped at a single locus. All of them could be assigned to an annotated gene. Thus even if a reference was available, our approach was able to retrieve *de novo* expressed variants of genes that were completely missed by the mapping.

**Figure 7:**
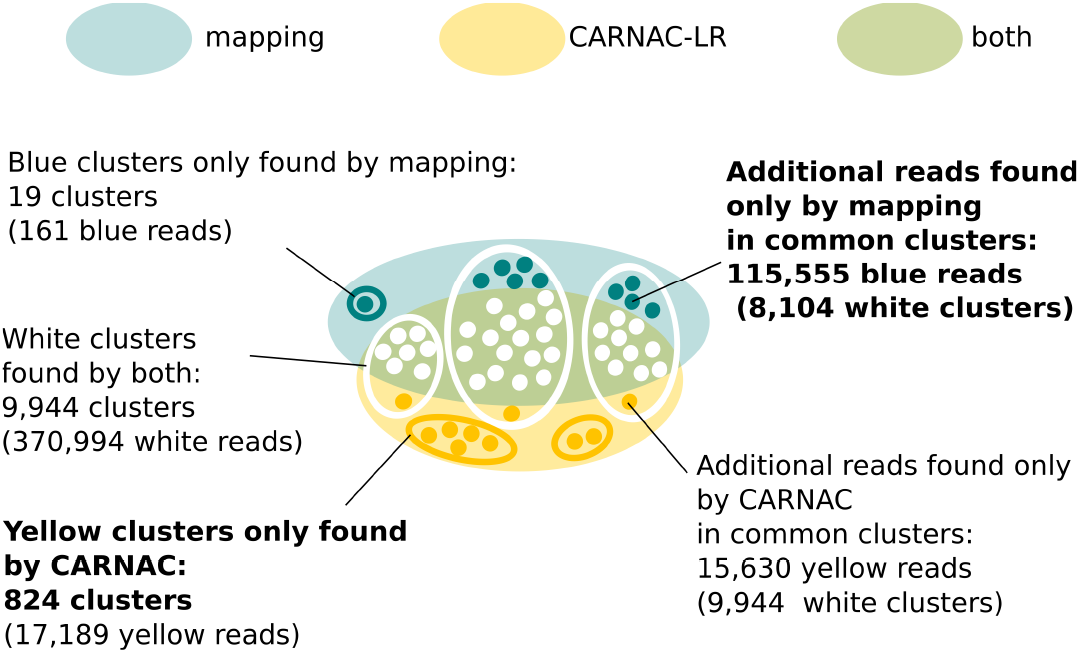
Complementarity of CARNAC-LR and mapping approaches. Only clusters of size 5 are represented. Mapping complemented common clusters a with a mean 13 reads per cluster in 90% of clusters. CARNAC-LR’s supply was tenfold lower with a mean 1,3 read added to 100% of common clusters. On the other hand, CARNAC-LR retrieved 15 fold more novel cluster than mapping.

##### Correlation of expression levels

Another way to look at these results is two consider the number of reads predicted by each method as the gene’s expression, and to compare expression levels predicted by our approach and by mapping. We shown that, despite the previously described differences, they are highly and linearly correlated, with a Pearson correlation coefficient of 0.80 (Figure 8).

**Figure 8:**
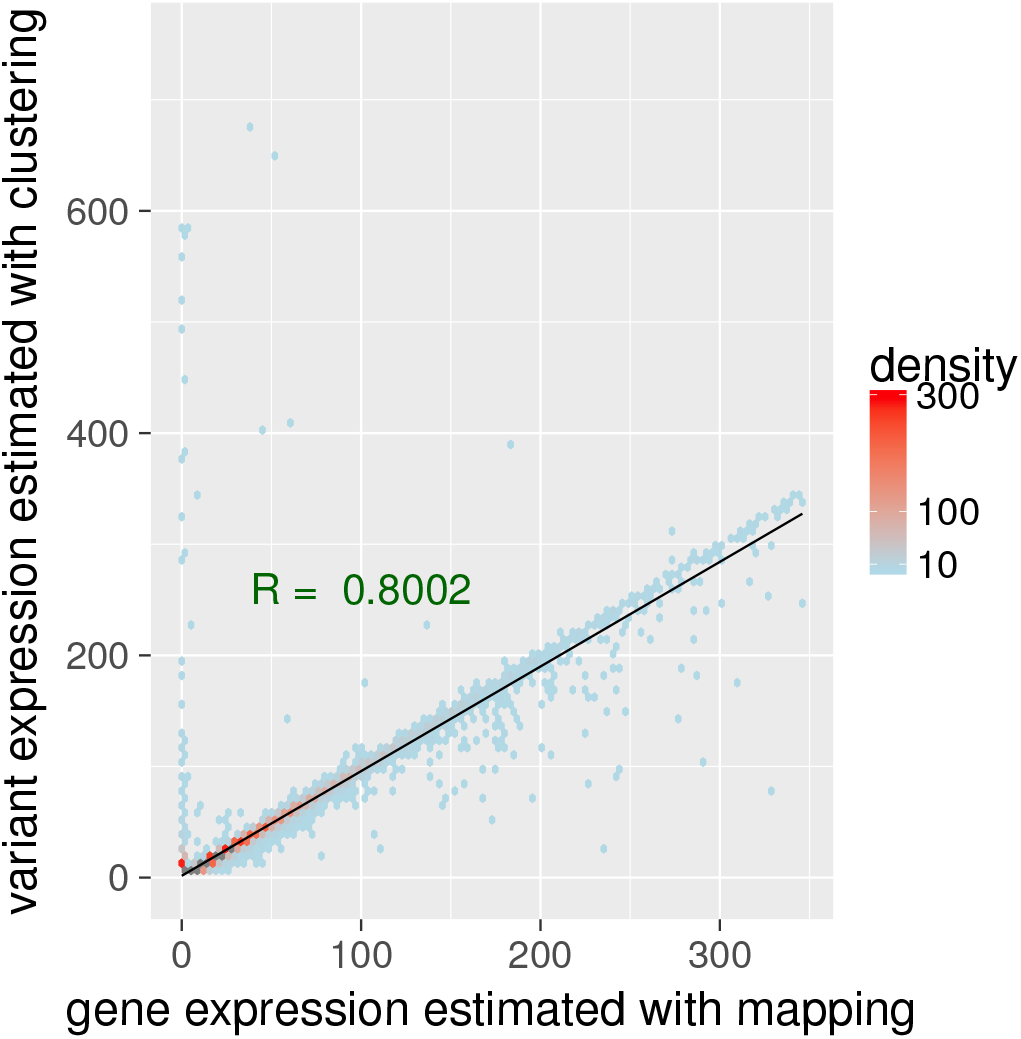
Comparison of clustering and mapping approaches. Comparison and correlation of expressions levels. Gene’s expression can be inferred by counting the number of reads by gene. For each gene we counted the number of reads retrieved by mapping and we compared it to the number of reads reported by our pipeline and validated by mapping. We computed the Pearson correlation coefficient between the two (in green). Density is the number of points counted in a colored region. Despite a few outliers, we can see a strong linear correlation between the two expression levels estimations (plotted in black). 7 outliers above 750 on Y axis (up to 3327) are not shown.

## 4 DISCUSSION

### 4.1 CARNAC-LR is well-suited on transcriptome instances

We demonstrated that our approach can compete with the state of the art algorithms to detect communities. On a rather small instance already, state of the art approaches show at least a lack of precision in comparison to our approach.

We showed that a *modularity*-based algorithms such as Louvain algorithm are not well-tailored for this problem, probably because of the heterogeneous size distribution of the clusters, and because of overlapping effects due to the repeats. Among tested state of the art approaches, only the CPM qualifies for retrieving clusters in our input graphs. However, by concentrating its results in a small subset of clusters, it obtains a poor recall and not all its predicted clusters can be trusted. On the other hand our approach shows a good consistency. We supplemented these results with a comparison with tools extensively used for clustering nucleic sequences, including developments used for EST clustering such as CD-HIT EST. We have shown that no published tool is currently capable of producing quality clusters from ONT RNA reads. We validated CARNAC-LR’s results using mouse transcriptome ONT reads, showing we can compute high confidence clusters for many genes. We highlight that the mapping procedure used for producing reference clusters for validation has its own limitations, thus the “ground truth” we refer to for the sake of clarity is in fact only partial.

### 4.2 CARNAC-LR can complement mapping approaches with respect to data with reference

Long reads enable to skip the transcript reconstruction step that is necessary with short reads, though difficult in particular when it involves assembly. Therefore, long reads constitute an interesting novel way to obtain reference transcripts. However, only a fraction of long reads are processed by mappers and downstream analysis is made difficult because of the error rates. In this context, our approach is shown to be an alternative approach to mapping for the identification of genes’ transcripts. We have shown that our pipeline could be a complementary procedure when reads can be mapped to a reference. Thus it tends to recover some clusters missed by mapping and allows a more efficient use of data. We have demonstrated a straightforward use case of our pipeline as a good proxy to access the expression levels by gene. ONT sequences have been shown to qualify for transcript quantification in [9]. In a long read sequencing experiment, it is likely that some reads contain too many errors to be mapped on a genome. CARNAC-LR can help identifying the origin gene of such reads, if they are put in cluster with other mapped reads. Moreover CARNAC-LR provides structured information that can be a sound input to other applications. For instance, a read correction step can be performed on each cluster instead of processing the whole data, in order to obtain high quality reference transcripts.

### 4.3 CARNAC-LR applies on non model species and ONT data

Non model species require *de novo* approaches, and two bioinformatics tools dedicated to them have emerged so far [49, 48]. Both comprise a pipeline conceived to process Pacific Biosciences Isoseq [3] reads only and require high quality long reads. Thus they could not be used on the data presented here. On the other hand CARNAC-LR is a generic approach that is designed to be used regardless of Third Generation Sequencing error profile and protocol. As a consequence it is the first method to perform *de novo* clustering on RNA reads from ONT.

### 4.4 Paralogy and repeats

The clustering of sequences from transcriptome reads is made difficult by the existence of multiple repeats. This first attempt to cluster RNA reads by gene is not designed to precisely assign reads from paralog genes to their original locus. We argue that particular instances such as paralog genes constitute research themes on their own and the clustering provides first-approximation results in these cases. We can think of a second clustering pass with the development of an adapted similarity calculation. CARNAC-LR gathers all reads from a gene family together, provided the different copies have not diverged too much and can thus be seen as a useful pre-processing step for the analysis of paralogs. A related research axis would be to describe how repeats like transposable elements that can be found in exons or retained introns are treated by the clustering procedure.

## 5 Conclusion

We propose a method designed for clustering long reads obtained from transcriptome sequencing in groups of expressed genes. New algorithmic challenge arises of the combination of a high error rate in data [7, 8], a high heterogeneity of coverage typical from expression data and an important volume of data. To this extent our question differs from EST clustering problems for instance. We demonstrated our method’s relevance for this application, in comparison to literature approaches. It takes reads early after their generation, without correction or filter. From the clusters, the expressed variants of each gene are obtained and related transcripts are identified, even when no reference is available. To make our solution practical for users, we provide an implementation called CARNAC-LR that, combined to Minimap, scales and is able to process quickly real data instances, as demonstrated by the processing of the whole mouse brain transcriptome.

As a consequence of the quick evolution of TGS, the sequencing field is frequently upgraded with new types of sequences. or instance, recent long read technology ONT RNA-direct could unlock amplification biases issues in RNA sequencing and thus is promising for gene expression studies (see Garalde et al., Highly parallel direct RNA sequencing on an array of nanopores, *bioRxiv*, 2016). But it shows higher error rates, at least comparatively to reads presented in this study, according to unpublished works. By proposing a generic tool that is tailored to these technologies, we wish to promote and encourage a broader use of long reads for transcriptome analysis.

### Data availability and Implementation

CARNAC-LR is written in C++, open source and available for Linux systems at github.com/kamimrcht/CARNAC-LR under the Affero GPL license.

The nanopore reads from the mouse RNA sample are available from the ENA repository under the following study: ERP107503.

